# Synaptotagmin 7 differentially regulates synaptic vesicle trafficking at tonic and phasic synapses

**DOI:** 10.64898/2026.11.09.765402

**Authors:** Michiko O. Inouye, Yulia Akbergenova, Zhuo Guan, Andrés B. Crane, Mónica C. Quiñones-Frías, J. Troy Littleton

## Abstract

Synaptic vesicle (SV) release probability can differ across neuronal subtypes, but the molecular mechanisms establishing such differences remain incompletely understood. In the current study we demonstrate that Synaptotagmin 7 (SYT7), a Ca²⁺-sensor regulating SV trafficking, is enriched at *Drosophila* tonic Ib synapses compared with phasic Is synapses. Loss of SYT7 preferentially enhanced evoked release at Ib terminals, whereas overexpression strongly suppressed fusion at both neuronal subtypes. High-frequency stimulation partially overcomes this inhibition, producing robust facilitation. Ultrastructural analysis revealed that SYT7 overexpression preserved morphologically docked SVs while promoting accumulation of enlarged presynaptic cisternae and redistribution of SV-associated proteins. Conversely, SYT7 loss reduced activity-induced presynaptic cisternae, suggesting an important role in regulating the bulk endocytosis pathway. Together, these findings identify differential SYT7 abundance as a contributor to tonic and phasic synaptic diversity and reveal a role for SYT7 in coordinating SV release and presynaptic membrane cycling.

**Summary:** Although *Drosophila* tonic and phasic motoneurons exhibit distinct neurotransmitter release properties, the underlying mechanisms remain unclear. We demonstrate that differential abundance of the Ca²⁺ sensor Synaptotagmin 7 contributes to their unique presynaptic output by regulating synaptic vesicle cycling.

## Introduction

Neuronal subtypes exhibit distinct morphologies, connectivity, excitability, and synaptic properties (Peng et al., 2021; Scala et al., 2021; Zeng, 2022). The *Drosophila* neuromuscular junction (NMJ) provides a useful model for identifying mechanisms underlying synaptic diversity, as larval muscles are co-innervated by tonic Ib and phasic Is motoneurons (MNs) with distinct release properties (Aponte-Santiago and Littleton, 2020). Tonic Ib synapses exhibit low synaptic vesicle (SV) release probability (*Pr*) and facilitation, whereas phasic Is synapses display higher *Pr* and depression (Newman et al., 2017; Lnenicka and Keshishian, 2000; Johansen et al., 1989). Although several proteins contributing to these differences have been identified (Jetti et al., 2023; Stark et al., 2026; Newman et al., 2022; Sauvola et al., 2021), how their unique synaptic output is established is largely unclear.

Synaptotagmins (SYTs) are a family of highly conserved proteins with dual Ca²⁺-binding C2A and C2B domains that regulate SV trafficking and fusion. SYT1 functions as the principal Ca²⁺ sensor for synchronous SV fusion, while SYT7 has been implicated in asynchronous release, short-term facilitation and SV recycling (Quiñones-Frías and Littleton, 2021). In *Drosophila*, SYT7 negatively regulates evoked SV release in a dosage-dependent manner, with loss of SYT7 enhancing release and overexpression suppressing fusion (Guan et al., 2020; Fujii et al., 2021). Differences in SYT7 abundance relative to SYT1 have been implicated in neuronal subtype-specific release properties in the mammalian cerebellum (Turecek and Regehr, 2019; Weyrer et al., 2021). Whether differential SYT7 abundance contributes more broadly to synaptic diversity in unknown.

Here, we demonstrate that SYT7 is enriched at *Drosophila* tonic Ib synapses compared to phasic Is synapses. Loss of SYT7 preferentially enhances evoked release at Ib terminals, whereas overexpression strongly suppresses spontaneous and evoked fusion at both synapse types. High-frequency stimulation partially overcomes this inhibition, producing robust facilitation of evoked responses. Despite severely reduced release, SYT7 overexpressing terminals retain morphologically docked SVs but show extensive redistribution of SV-associated proteins and accumulation of enlarged presynaptic cisternae. In contrast, loss of SYT7 reduces activity-dependent cisternae formation associated with bulk endocytosis. Together, these findings identify differential SYT7 abundance as a mechanism contributing to tonic and phasic synaptic output differences and reveal an additional role for SYT7 in presynaptic membrane trafficking.

## Results

### SYT7 is enriched at tonic Ib synapses and preferentially suppresses their release probability

We previously found that *Drosophila* SYT7 inhibits evoked SV release in a dosage-dependent manner (Guan et al., 2020). To determine whether differential SYT7 abundance may contribute to the distinct output of tonic Ib and phasic Is MNs, we examined the levels of endogenously GFP-tagged SYT7 at larval NMJs. SYT7-GFP fluorescence was ∼2.1-fold higher at Ib than Is terminals across muscles 6/7, 4, and 1 (Fig. 1A–E). In contrast, the SV Ca²⁺ sensor SYT1 showed a smaller ∼1.3-fold Ib enrichment (Fig. 1F–H). Accordingly, the SYT7 Ib/Is abundance ratio was higher than SYT1 across all NMJs (Fig. 1I, J). These data indicate SYT7 is differentially enriched at presynaptic terminals between the two MN subtypes (Fig. 1K).

**Figure 1.**
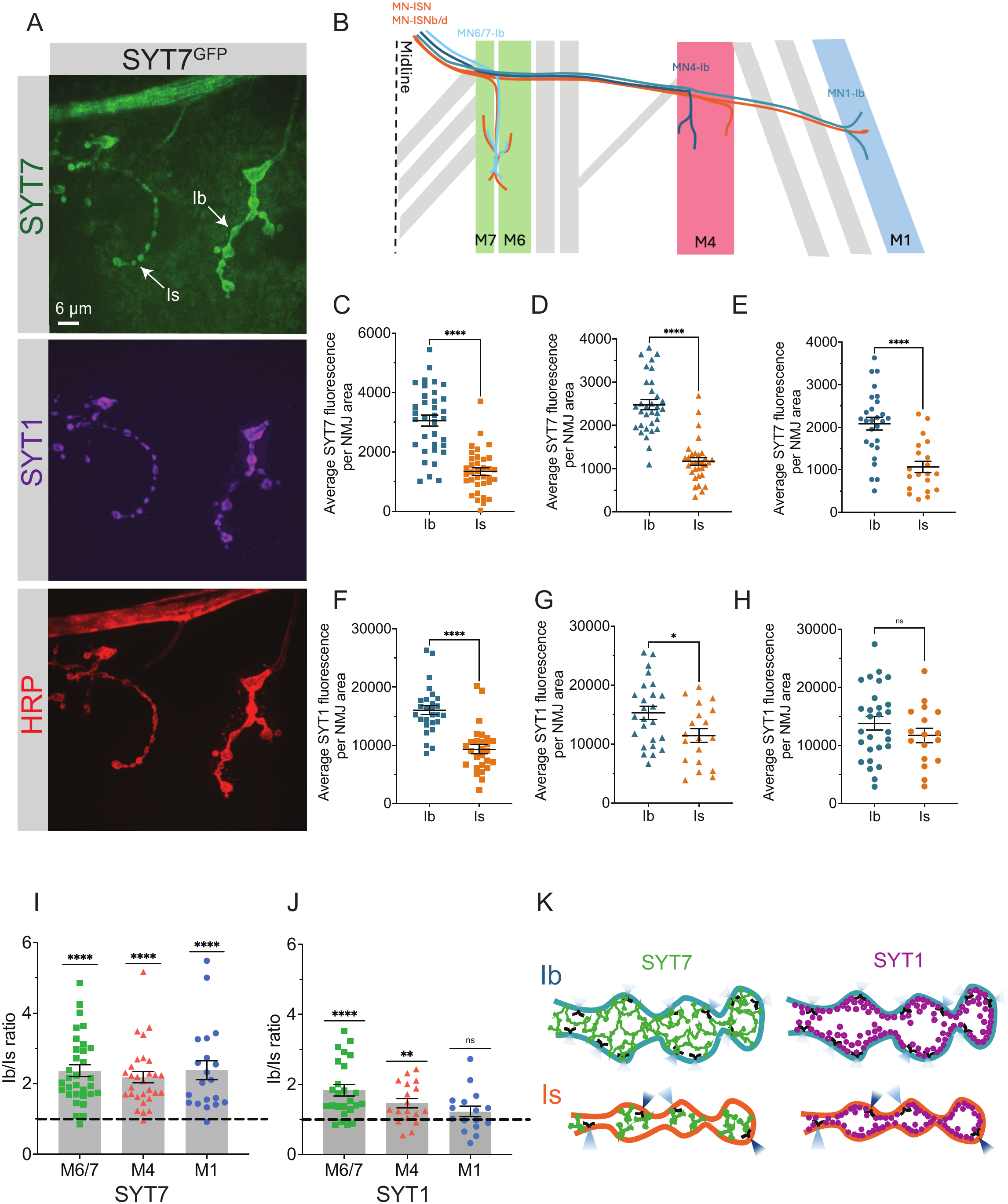
SYT7 expression levels are ∼2 fold higher at Ib NMJs. **(A)** Immunocytochemistry of 3^rd^ instar muscle 4 NMJs with anti-HRP (red) and anti-SYT1 (magenta) in CRISPR-tagged SYT7^GFP^ (green) larvae. Ib and Is terminals are indicated with arrows. Scale bar = 6 μm. (**B)** Schematic representing Ib and Is innervation of a larval abdominal hemisegment with muscles (M) 6/7, 4, and 1 indicated. Quantification of SYT7^GFP^ **(C-E)** and SYT1 **(F-H)** expression in Ib and Is terminals at muscles 6/7, 4 and 1. Quantification of relative ratios of **(I)** SYT7 and **(J)** SYT1 protein levels in Ib/Is within the same NMJ for muscles 6/7, 4, and 1. **(K)** Diagram highlighting higher SYT7 protein level versus SYT1 in Ib terminals compared to Is. SYT7 localization (green) is shown in its internal compartment and at the plasma membrane near AZ T-bars (black). SYT1 is localized to SVs (magenta). Blue cones indicate differential AZ *P_r_*, with darker colors representing higher *P_r_*. Data is shown as mean ± SEM. ** p<0.01; *** p<0.001; **** p<0.0001; ns, not significant.

To test whether this difference contributes to their unique release properties, we mapped evoked *P_r_* at individual active zones (AZs) using optical quantal imaging. Consistent with prior observations, control Is AZs exhibited higher average *P_r_* than Ib AZs (Fig. 2A, B). In *Syt7^M1^* null mutants, average *P_r_* increased 2.3-fold at Ib AZs but was not significantly altered at Is AZs (Fig. 2A–C). Thus, endogenous SYT7 preferentially limits evoked release at tonic Ib synapses. We next examined activity-dependent membrane uptake and recycling using the lipophilic FM1-43 dye (Fig. 2D-F). Following 10 Hz stimulation, control Is terminals accumulated significantly more dye than Ib terminals as expected. *Syt7^M1^*nulls showed a ∼1.5-fold increase in FM1-43 loading at Ib terminals but only a ∼1.1-fold change at Is synapses (Fig. 2D, E). Dye unloading with 10 Hz stimulation was also preferentially enhanced at Ib NMJs in *Syt7^M1^*, consistent with increased SV release at Ib terminals (Fig. 2D, F). Together, quantal imaging and FM1-43 uptake and release indicate higher levels of endogenous SYT7 expression at Ib terminals contributes to their lower AZ *P_r_* and decreased evoked output (Fig. 2G).

**Figure 2.**
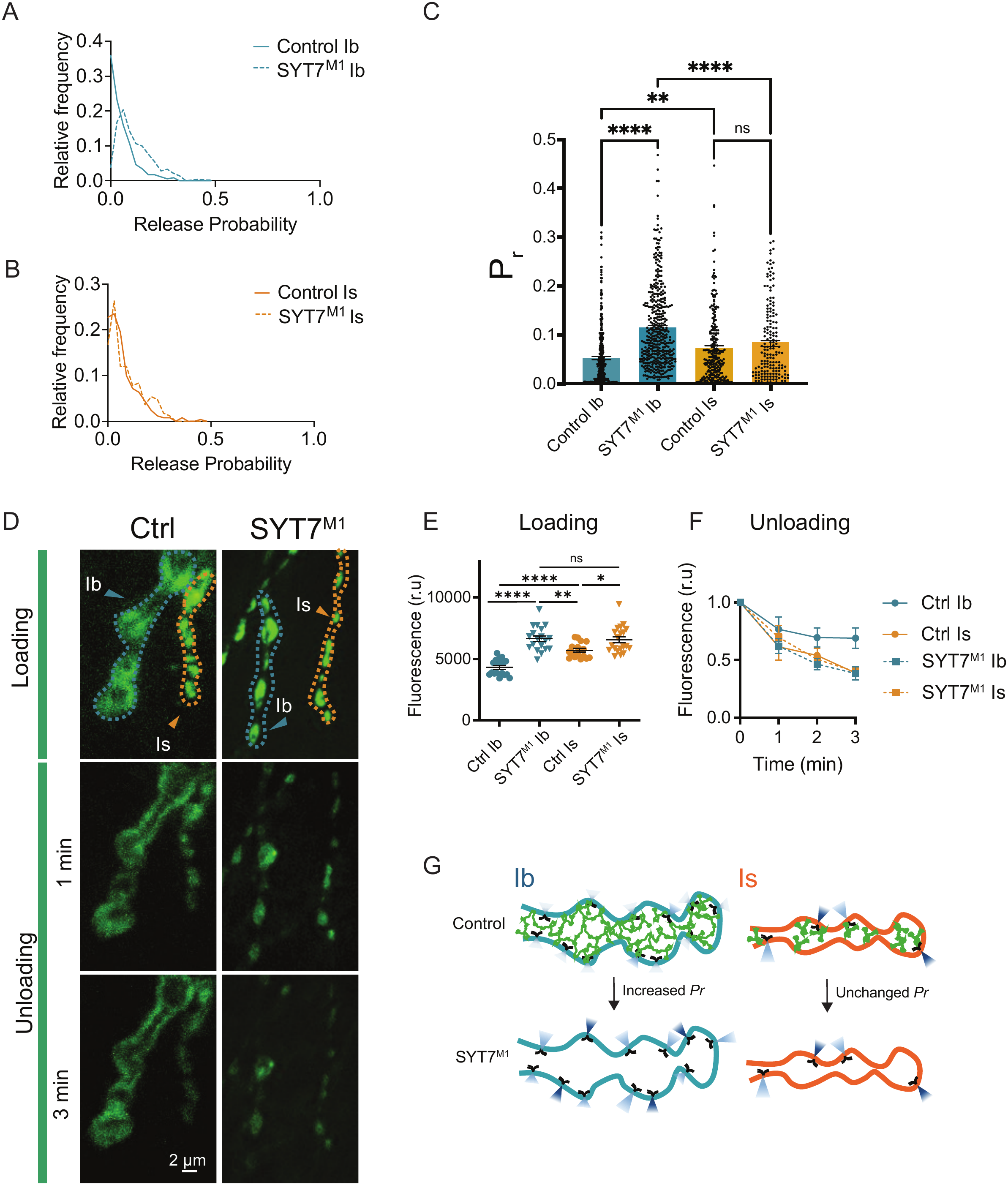
Loss of SYT7 preferentially increases *P_r_* at Ib NMJs. **(A)** *Pr* frequency distributions for control (cac^RFP^;;Lex-GCaMP7s, GluRIIB-GFP/TM3, solid line) and SYT7^M1^ (cac^RFP^;;Lex-GCaMP7s, GluRIIB-GFP/TM3; SYT7^M1^, dashed line) at muscle 4 Ib NMJs. **(B)** *Pr* frequency distributions for control (solid line) and SYT7^M1^ (dashed line) at muscle 4 Is NMJs. **(C)** Plot showing mean *P_r_* for all individual AZs measured per condition in panels A and B. **(D)** Representative fluorescence images of FM1-43 dye loading and unloading in control and *Syt7^M1^* NMJs. Loading was performed for 2 min at 10 Hz stimulation, followed by unloading at 10 Hz and imaged at 1 min intervals. Type Ib and Is branches are indicated by cyan and orange arrowheads and dashed outlines, respectively. Scale bar = 2 µm. **(E)** Quantification of fluorescence intensity following loading reveals higher dye uptake in Is versus Ib boutons in control NMJs. In contrast, Syt7^M1^ mutants exhibit comparable intensities in Ib and Is boutons due to increased loading in Ib terminals. Control FM1-43 levels: Ib (4310 ± 153.3, 18 NMJs, 5 larvae); Is (5703 ± 148.2, 17 NMJs, 4 larvae). *Syt7^M1^* Ib FM1-43 levels: (6645 ± 227.9, 19 NMJs, 5 larvae); Is (6548 ± 242.7, 20 NMJs, 5 larvae). **F)** Quantification of dye unloading during continuous 10 Hz stimulation for 3 min. Fluorescence intensity was normalized to the initial fluorescence measured immediately after loading (time 0, set to 1 for each NMJ). Normalized fluorescence was measured after 1, 2, and 3 min of stimulation for Ib and Is boutons in control and *Syt7^M1^* mutants. *Syt7^M1^* Ib terminals exhibit a more pronounced decrease in fluorescence than control Ib boutons, whereas Is terminals show similar unloading kinetics in both genotypes. **(G)** Model showing that loss of SYT7 in specifically increased release in Ib terminals, whereas Is remains unaltered. Data = mean ± SEM. Statistical analysis was performed with one-way ANOVA with Tukey correction. **** p<0.0001; ns, not significant.

### SYT7 overexpression inhibits neurotransmitter release at both tonic and phasic synapses

To determine how elevated SYT7 abundance affects Ib and Is MNs, two UAS-SYT7 transgenic lines were generated. Expression of UAS-SYT7^OE1^ and UAS-SYT7^OE2^ transgenes with the pan-MN driver OK6-GAL4 increased SYT7 protein abundance by ∼2-fold and 3.4-fold, respectively, in adult head extracts compared to controls (Fig. S1A, B). Immunostaining with anti-SYT7 antisera showed OK6-GAL4 driven overexpression eliminated the endogenous difference in SYT7 abundance between Ib and Is terminals in both lines, although SYT7^OE2^ exhibited higher overall expression compared to SYT7^OE1^ (Fig. S1C, D). SYT7 immunoreactivity was distributed throughout synaptic boutons in these lines, including at the peri-AZ and within internal regions of the bouton, along with some signal at the plasma membrane near BRP-positive AZs (Fig. S1E). These locations position SYT7 to regulate both SV exocytosis and recycling pathways.

To determine how SYT7 overexpression alters presynaptic output, FM1-43 uptake during 10 Hz stimulation was quantified. Whereas control Is terminals accumulated more dye than Ib, SYT7 overexpression markedly reduced loading at both Ib and Is NMJs (Fig. 3A, B). Optical quantal imaging revealed dramatic reductions in AZ *P_r_* at both Ib and Is terminals following SYT7 overexpression (Fig. 3C–E). The proportional reduction in evoked release was greater at Is AZs, demonstrating the phasic fusion machinery remains highly sensitive to elevated SYT7 levels despite its lower endogenous abundance. Two-electrode voltage-clamp recordings confirmed a strong, dose-dependent reduction in evoked currents in SYT7^OE1^ and SYT7^OE2^ larvae (Fig. 3F–H). Despite strong suppression of synchronous release, normalized current traces revealed enhanced asynchronous fusion after stimulation (Fig. 3I, J). SYT7 overexpression also increased failure rates following stimulation in 2 mM extracellular Ca²⁺, from 0% in controls to 7.3% in SYT7^OE1^ and 54.3% in SYT7^OE2^ (Fig. 3K). Spontaneous release frequency was also dramatically reduced, while miniature amplitude was unchanged (Fig. 3L, M). We conclude that increasing SYT7 abundance strongly suppresses evoked *P_r_* and spontaneous SV fusion at both Ib and Is synapses (Fig. 3N).

**Figure 3.**
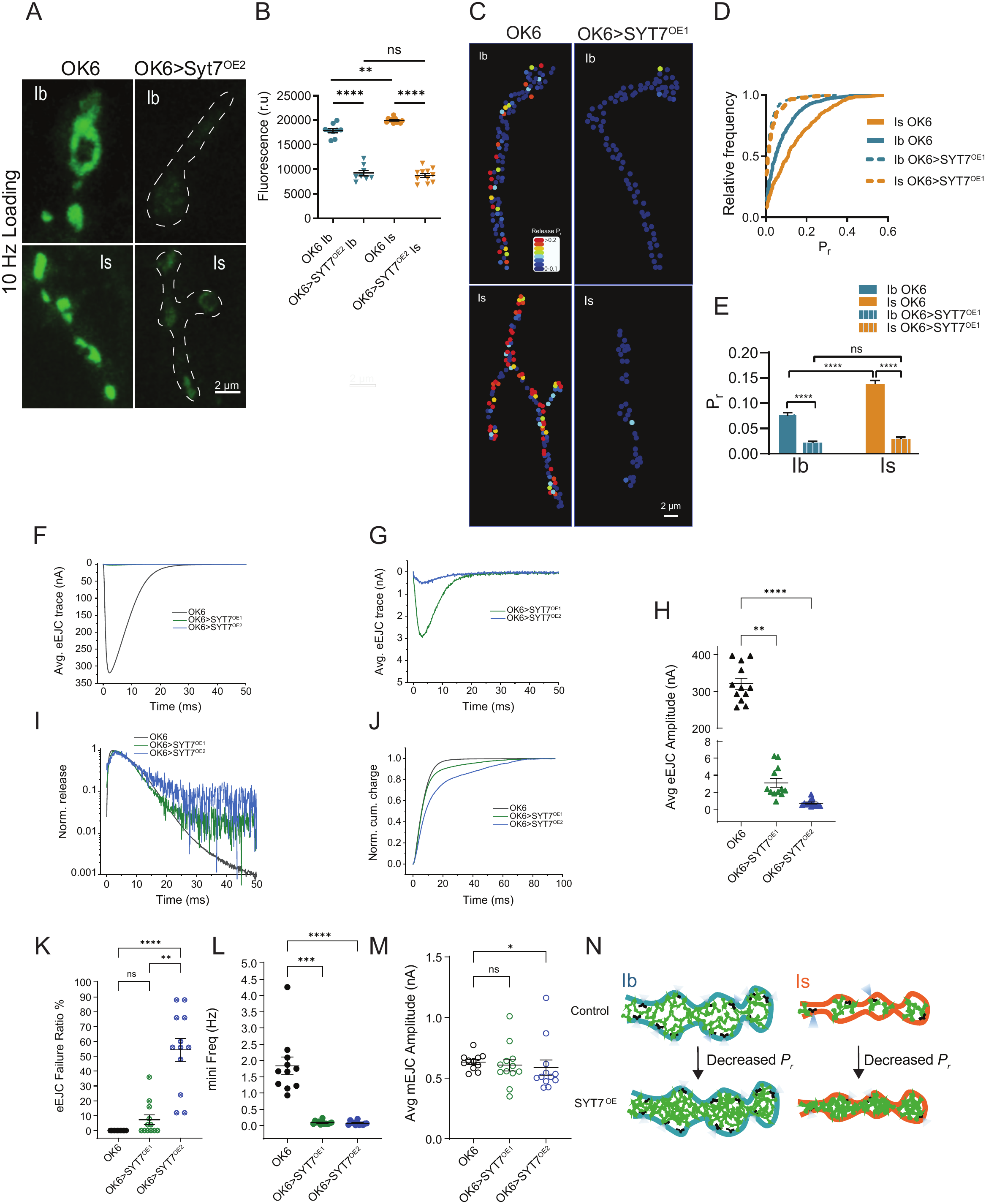
SYT7 overexpression reduces *Pr* at both Ib and Is NMJs. **(A)** Representative images of FM1-43 loading at type Ib and Is boutons of control (OK6-GAL4) and SYT7-overexpressing (OK6>SYT7^OE2^) NMJs following 5 min stimulation at 10 Hz. Dashed outlines highlight boutons showing reduced FM1-43 fluorescence in SYT7-overexpressing NMJs. Scale bar = 2 µm. **(B)** Quantification of FM1-43 fluorescence intensity demonstrates a significant reduction in dye loading at both Ib and Is boutons following SYT7 overexpression. In controls, Is boutons exhibit significantly higher dye uptake than Ib boutons, whereas this difference is abolished following SYT7 overexpression. Fluorescence levels for OK6 Ib boutons: (17857 ± 1326, 9 NMJs, 4 larvae); OK6 Is boutons (19940 ± 485.5, 12 NMJs, 4 larvae); OK6>SYT7^OE2^ Ib boutons: (9216 ± 1635, 8 NMJs, 4 larvae); OK6>SYT7^OE2^ Is boutons (8737 ± 1397, 11 NMJs, 4 larvae). **(C)** Representative AZ evoked *P_r_* heatmaps for Ib and Is boutons in control (OK6-GAL4) and SYT7-overexpressing (OK6>SYT7^OE1^) 3^rd^ instar larvae for 1 Hz nerve stimulation. Scale bar = 2 µm. **(D)** Cumulative frequency distributions of individual AZ *P_r_* in Is and Ib boutons from control and SYT7-overexpressing NMJs. Under control conditions, Is AZs exhibit higher *P_r_* than Ib, as indicated by the rightward shift in their cumulative distribution. SYT7 overexpression markedly shifts both distributions toward lower *P_r_*, eliminating the characteristic difference between Is and Ib AZs. **(E)** Quantification of mean AZ *P_r_* in Is and Ib boutons. Average release *Pᵣ*: control Ib AZs: (0.07811 ± 0.0032, 6 NMJs, 6 larvae); control Is AZs: (0.1386 ± 0.006, 5 NMJs, 5 larvae); OK6>SYT7^OE1^ Ib AZs: (0.02263 ± 0.0017, 6 NMJs, 6 larvae); OK6>SYT7^OE1^ Is AZs: (0.029 ± 0.0035, 7 NMJs, 7 larvae). **(F-G)** Average eEJC traces recorded from OK6 (black), OK6>SYT7^OE1^ (green), and OK6>SYT7^OE2^ (blue) 3^rd^ instar larval NMJs. Average eEJC traces were obtained by summing all eEJC events under the first peak distribution. Note the different scale bars in F and G. **(H)** Quantification of mean eEJC amplitudes in the indicated genotypes (OK6: 320.6 ± 14.8 nA, n=12; OK6>SYT7^OE1^: 3.09 ± 0.5 nA, n=12; OK6>SYT7^OE2^: 0.7 ± 0.1 nA, n=12). Statistical analysis was performed with ordinary one-way ANOVA test with Dunnett’s multiple comparisons test. **(I)** Average normalized responses for each genotype are plotted on a semi-logarithmic graph to display release components. **(J)** Cumulative release normalized for the maximum in 2.0 mM Ca^2+^ for each genotype. Each trace was adjusted to a double exponential fit. **(K)** Quantification of eEJC failure ratio (%) in the indicated genotypes (OK6: 0 ± 0, n=12; OK6>SYT7^OE1^: 7.33 ± 3.26, n=12; OK6>SYT7^OE2^: 54.33 ± 7.68, n=12). Statistical analysis was performed with one-way Anova Sidak’s multiple comparisons test. **(L)** Quantification of mean mEJC frequency in the indicated genotypes (OK6: 1.84 ± 0.27 Hz, n=11; OK6>SYT7^OE1^: 0.092 ± 0.016 Hz, n=12; OK6>SYT7^OE2^: 0.070 ± 0.018 Hz, n=12). Statistical analysis was performed with ordinary one-way ANOVA test with Dunnett’s multiple comparisons test. **(M)** Quantification of mean mEJC amplitude in the indicated genotypes (OK6: 0.63 ± 0.02 nA, n=11; OK6>SYT7^OE1^: 0.61 ± 0.05 nA, n=12; OK6>SYT7^OE2^: 0.59 ± 0.06 nA, n=12). Statistical analysis was performed with ordinary one-way ANOVA test with Dunnett’s multiple comparisons test. **N)** Model showing decreased *Pr* as a result of SYT7 overexpression in both Ib and Is NMJs. Data = mean ± SEM. * p<0.05; ** p<0.01; *** p<0.001; **** p<0.0001; ns, not significant.

Consistent with functional defects in presynaptic output, SYT7 overexpression caused dose-dependent developmental defects, including impaired crawling, reduced pupation height, and pharate adult lethality (Fig. S1F-I). Rare escaper adults showed impaired climbing and defects in wing expansion, phenotypes consistent with reduced synaptic transmission and subsequent muscle weakness (Fig. S1G). SYT7 overexpression also caused mild NMJ morphological defects, including larger synaptic boutons, fewer boutons and a small reduction in AZ number (Fig. S1J-N). Enlarged HRP-positive inclusions were also observed within synaptic boutons and nerve bundles (Fig. S1J, O, Fig. S2). Large vacuolar structures in synapses have been reported previously following SYT7 overexpression in mammals (Virmani et al., 2003) and may represent internal compartments that retain SVs and restrict their availability for fusion (see below). These observations suggest broader consequences for excessive SYT7 expression on membrane trafficking beyond the strong reduction in SV fusion.

### High-frequency stimulation partially overcomes SYT7-mediated suppression of SV fusion

Given SYT7’s ability to bind Ca^2+^ and its role in activity-induced facilitation, changes in SYT7 function during high frequency firing and elevated intra-terminal [Ca^2+^] are likely to modulate its interaction with the SV fusion machinery. The marked reduction in both evoked and spontaneous SV release raises the question of whether SYT7 overexpression constitutively blocks SV release or leaves a population capable of enhanced fusion during sustained activity. To test these possibilities, evoked responses were recorded during a stimulation paradigm of 30 stimuli at 0.5 Hz, followed by 500 stimuli at 10 Hz, and a return to 0.5 Hz stimulation. At low frequency, both overexpression lines greatly reduced evoked responses relative to controls. During the 10 Hz train, control NMJs displayed marked depression as readily-releasable SV pools were depleted, while SYT7 overexpressing terminals showed a robust increase in evoked responses (Fig. 4A–C). The increase evoked response was particularly evident at the transition to 10 Hz, when responses that had been strongly suppressed at 0.5 Hz rapidly facilitated (Fig. 4D, E). Although both SYT7^OE1^ and SYT7^OE2^ larvae displayed facilitation, the absolute evoked current amplitude remained well below control amplitudes throughout the 10 Hz train (Fig. 4A–C). Following the return to 0.5 Hz, evoked release rapidly increased in controls by the 2^nd^ stimuli, while it was strongly suppressed to the prior reduced output level in overexpression lines (Fig. 4F, G). These data indicate SYT7 normally clamps release at low intracellular Ca^2+^ levels in a dose-dependent manner. The rise in intracellular Ca^2+^ during 10 Hz stimulation releases this brake to transiently increase evoked *P_r_*, with the clamp being rapidly restored once Ca^2+^ levels fall.

**Figure 4.**
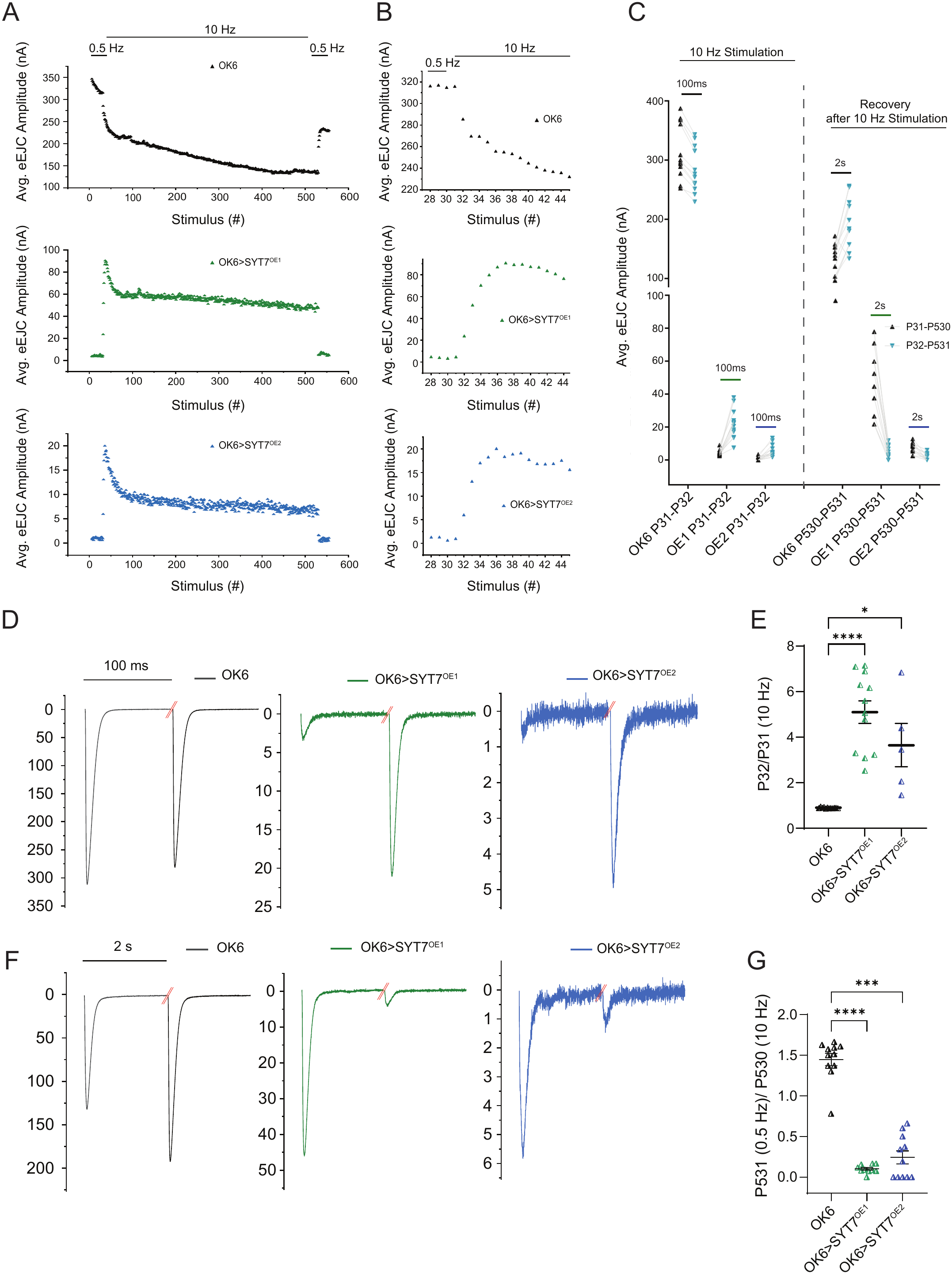
Evoked responses during and after short-term stimulation in SYT7 overexpression lines. **(A)** Average eEJC amplitudes for the indicated stimulation protocol in 2 mM external Ca^2+^ for OK6 (black), OK6>SYT7^OE1^ (green), and OK6>SYT7^OE2^ (blue). **(B)** Representative average eEJC amplitudes for the last 3 responses to 0.5 Hz and the 1^st^ 14 responses to 10 Hz stimulation are shown. **(C)** Quantification of mean eEJC amplitudes for the 1^st^ eEJC (P31) and the 2^nd^ eEJC (P32) at the onset of 10 Hz stimulation for the indicated genotypes (P31 in OK6: 313.46 ± 13.38, n = 12; P32 in OK6: 282.97 ± 11.01, n = 12; P31 in OK6>SYT7^OE1^: 4.43 ± 0.50, n = 12; P32 in OK6>SYT7^OE1^: 21.89 ± 2.60, n = 12; P31 in OK6>SYT7^OE2^: 0.91 ± 0.35, n = 12; P32 in OK6>SYT7^OE2^: 5.95 ± 1.10, n = 12). Quantification of mean eEJC amplitudes for the last eEJC (P530) response to 10 Hz and the 1^st^ eEJC (P531) response to 0.5 Hz for the indicated genotypes (P530 in OK6: 134.43 ± 7.58, n = 11; P531 in OK6: 192.23 ± 13.07, n = 11; P530 in OK6>SYT7^OE1^: 47.35 ± 6.40, n = 10; P531 in OK6>SYT7^OE1^: 5.27 ± 1.18, n = 10; P530 in OK6>SYT7^OE2^: 7.50 ± 0.98, n = 11; P531 in OK6>SYT7^OE2^: 1.78 ± 0.61, n = 11). **(D)** Representative eEJC traces of P31 and P32 for OK6 (black), OK6>SYT7^OE1^ (green), and OK6>SYT7^OE2^ (blue). The time interval between the stimuli is 100 ms. **(E)** Quantification of facilitation (P32/P31 ratio) of eEJC amplitudes in the indicated genotypes (OK6: 0.90 ± 0.007, n=12; OK6>SYT7^OE1^: 5.10 ± 0.49, n=12; OK6>SYT7^OE2^: 3.64 ± 0.95, n=5). **(F)** Representative eEJC traces of responses during the transition from 10 Hz (P530) to 0.5 Hz (P531) stimulation for OK6 (black), OK6>SYT7^OE1^ (green), and OK6>SYT7^OE2^ (blue). The interval time between the stimuli is 2 sec. **(G)** Quantification of the P531/P530 ratio by eEJC amplitude in the indicated genotypes (OK6: 1.45 ± 0.08, n=11; OK6>SYT7^OE1^: 0.10 ± 0.016, n=10; OK6>SYT7^OE2^: 0.24 ± 0.08, n=11). Data = mean ± SEM. Statistical analysis was performed with ordinary one-way ANOVA test with Dunnett’s multiple comparisons test: *** p<0.001; **** p<0.0001.

### Overexpressed SYT7 redistributes SV-associated proteins within presynaptic boutons

Prior studies demonstrated SYT7 normally slows SV entry into the readily releasable pool (RRP) (Guan et al., 2020), suggesting SYT7 overexpression may alter additional aspects of presynaptic membrane cycling and SV organization. To test this possibility, FM1-43 dye uptake studies were performed following 100 Hz stimulation to analyze SV cycling. 100 Hz stimulation resulted in robust dye uptake in SYT7 overexpression lines (Fig 5A, B). Although total FM-143 fluorescence was unchanged, SYT7 overexpression altered dye distribution, promoting accumulation in prominent internal puncta rather than the more uniform peripheral pattern observed in controls (Fig. 5A, C). To determine whether SV-associated proteins were similarly redistributed, we examined SYT1 and Complexin (CPX) immunoreactivity. In controls, both proteins exhibited peripheral enrichment consistent with SV localization near the plasma membrane. SYT7 overexpression disrupted this pattern, resulting in an irregular distribution and increased internal punctate accumulation (Fig. 5D–G). Quantification of peripheral versus central fluorescence, as well as intensity profiles across individual boutons, confirmed a redistribution of SV proteins from the periphery to internal compartments (Fig. 5D–I). Despite the striking changes in spatial distribution, total bouton-associated SYT1 and CPX fluorescence changed only modestly, and substantial signal remained near BRP-positive AZs (Fig. 5J–O). These observations indicate increased SYT7 levels alters the organization of SV-associated proteins within presynaptic terminals. Enlarged HRP-positive inclusions and changes in the distribution of Rab11, Rab5, and the endolysosomal marker NHE3 further indicate excess SYT7 affects multiple presynaptic membrane compartments (Fig. S2A-C). In contrast, mitochondrial distribution at NMJs was unaffected following SYT7 overexpression (Fig. S2D). We conclude that SYT7 overexpression alters endocytosis pathways, generating abnormal distributions of SV proteins within presynaptic terminals.

**Figure 5.**
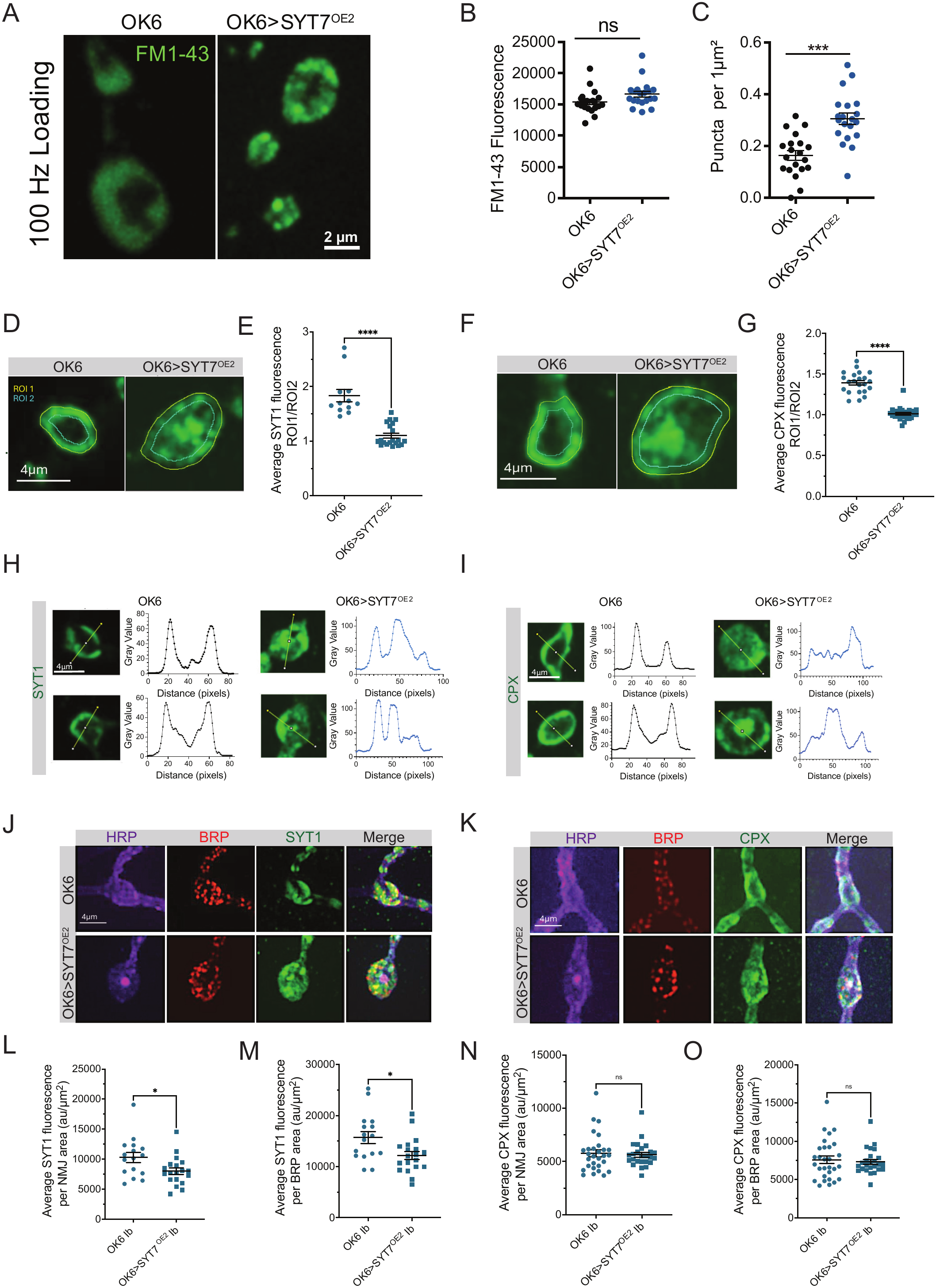
SYT7 overexpression results in altered SV distribution. **(A)** Representative images of FM1-43 loading at NMJs in control (OK6) and SYT7-overexpressing (OK6>SYT7^OE2^) boutons. FM1-43 loading was induced by high-frequency stimulation (100 Hz for 30 s). While control boutons exhibit a relatively uniform distribution of FM1-43 fluorescence, SYT7-overexpressing boutons display a prominent pattern of punctate fluorescence. Scale bar = 2 µm. **(B)** Quantification of mean FM1-43 fluorescence intensity per bouton reveals no significant difference between control and SYT7-overexpressing NMJs (ns, not significant). **(C)** Quantification of fluorescent puncta density (puncta/µm²) demonstrates a significant increase in discrete FM1-43-positive structures in SYT7-overexpressing boutons compared with controls (***p < 0.001). SYT1 **(D-E)** and CPX **(F-G)** immunolabeling in muscle 4 synaptic boutons of controls and OK6>SYT7^OE2^ larvae with ROIs tracing bouton periphery and inner boutons. The intensity ratio between the 2 ROIs are quantified on the right. SYT1 **(H)** and CPX **(I)** line density profile plots for distribution analyses are shown. Representative images of HRP, BRP, and SYT1 **(J)** and HRP, BRP, and CPX **(K)** immunolabeling with quantification of mean fluorescence per HRP area and mean fluorescence per BRP area for SYT1 **(L-M)** and CPX **(N-O)** Cpx. Data = mean ± SEM. Statistical analysis was performed with one-way ANOVA with Tukey correction and unpaired t-test. * p<0.05; *** p<0.001; ns, not significant.

### SYT7 overexpression increases presynaptic cisternae while preserving morphologically docked SVs

To examine membrane structures associated with these changes, transmission electron microscopy (TEM) was performed on Ib and Is boutons (Fig. 6A). In the absence of stimulation, control terminals contained abundant small SVs (Fig. 6A). SYT7-overexpressing Ib and Is boutons contained numerous enlarged membrane-bound cisternae in both MN subtypes, including structures greater than 50 nm in diameter (Fig. 6A–C). The accumulation of these larger internal membrane compartments is consistent with the puncta observed by FM1-43 loading and SV protein immunostaining. The number of morphologically docked SVs contacting the presynaptic membrane near AZ T-bars was not significantly altered by SYT7 overexpression in either Ib or Is terminals (Fig. 6D, E). The number of SVs within 200 nm of the T-bar showed only modest changes, suggesting SVs docking and distribution at AZs is relatively unaffected by SYT7 overexpression despite the profound loss of evoked release. These observations suggest SYT7 overexpression reduces SV fusogenicity or impairs SV priming downstream of morphological docking.

**Figure 6.**
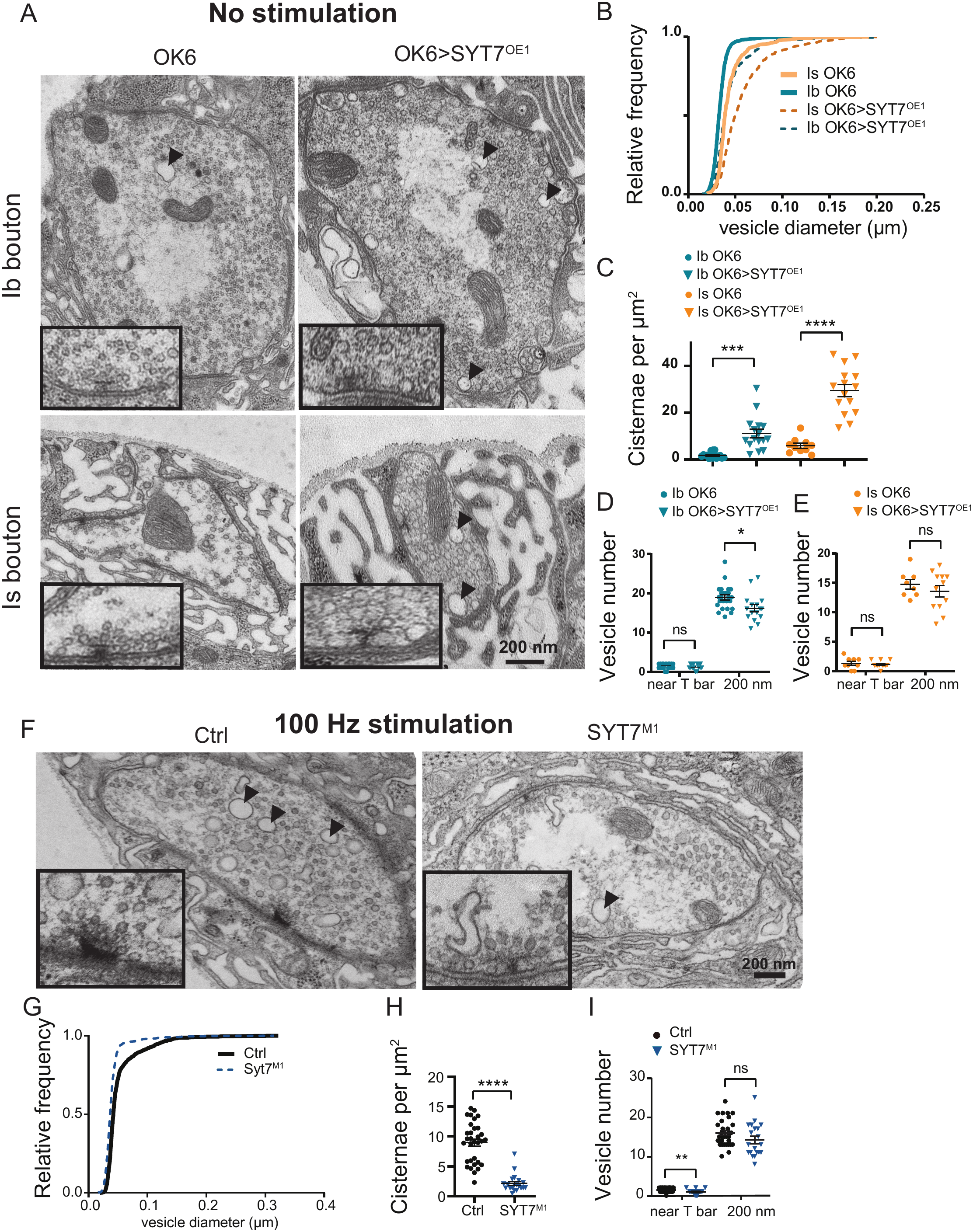
SYT7 promotes bulk endosome formation. **(A)** Representative TEMs of Ib and Is boutons from control (OK6) and SYT7-overexpressing (OK6>SYT7^OE1^) larvae fixed under resting conditions (no stimulation). Insets show higher-magnification views of SVs near AZs. Black arrows denote several cisternae in boutons for control or OK6>SYT7^OE1^. **(B)** Cumulative frequency distributions of vesicle diameters in control and SYT7-overexpressing Ib and Is boutons. Vesicle diameters were determined by drawing and measuring a line profile across individual vesicles in electron micrographs. Note the shift in the distribution toward larger vesicle sizes observed in boutons from SYT7-overexpressing larvae. **(C)** Quantification of the number of cisternae larger than 50 nm in diameter per μm² of bouton area in resting control and SYT7-overexpressing Ib and Is boutons. SYT7 overexpression significantly increases cisternae density in both bouton types. Each data point represent quantification from individual bouton micrographs. **(D, E)** Quantification of SV number immediately adjacent to the AZ T-bar and within a 200 nm radius of the AZ T-bar in Ib **(D)** and Is **(E)** boutons. Each data point represent quantification from an individual AZ in a bouton micrograph. **(F)** Representative TEMs of control and *Syt7^M1^* null mutant boutons fixed immediately following 100 Hz stimulation. Insets show higher-magnification views of the AZ and surrounding vesicle population. Black arrows denote several cisternae in boutons for control or *Syt7^M1^*. **(G)** Cumulative frequency distributions of vesicle diameters in control and *Syt7^M1^* null boutons following 100 Hz stimulation. **(H)** Quantification of cisternae larger than 100 nm per μm² of bouton area following 100 Hz stimulation for 30 sec. *Syt7^M1^* null boutons exhibit a significant reduction in cisternae density compared to controls. Data points represent individual analyzed boutons. **(I)** Quantification of SV numbers immediately adjacent to the T-bar and within a 200 nm radius of the T-bar following 100 Hz stimulation. *Syt7^M1^* mutant boutons show a significant reduction in vesicles near the T-bar, whereas vesicle numbers within 200 nm are not significantly different from controls. Individual data points represent analyzed AZs. Scale bar = 200 nm. ns, not significant; *p < 0.05, **p < 0.01, ***p < 0.001, ****p< 0.0001.

As enlarged cisternae can arise through the activity-dependent bulk endocytosis pathway (Akbergenova and Bykhovskaia, 2010; Clayton et al., 2008; Stevens et al., 2012; Vijayakrishnan et al., 2009), we assayed whether endogenous SYT7 is required for this mechanism. Following 100 Hz stimulation, control terminals contained numerous large cisternae as previously described for bulk endocytosis. In contrast, *Syt7^M1^* null boutons exhibited a marked reduction in these structures (Fig. 6F–H). Stimulated *Syt7^M1^* terminals also showed a reduction in SVs immediately adjacent to the T-bar (Fig. 6I), consistent with enhanced fusion in the absence of SYT7 (Fig. 2). The SV population within 200 nm of the T-bar was not significantly changed (Fig. 6I). Together, these experiments indicate SYT7 regulates the formation of large presynaptic membrane cisternae associated with bulk endocytosis. SYT7 overexpression promotes their accumulation or retention even at rest, whereas SYT7 loss reduces their formation or stability following strong stimulation.

## Discussion

The distinct release properties of *Drosophila* tonic Ib and phasic Is MNs are thought to arise in part from differences in presynaptic organization and the molecular machinery controlling SV release (Sauvola et al., 2021; Jetti et al., 2023; Bakshinska et al., 2025; He et al., 2023; Stark et al., 2026; Medeiros et al., 2024; Newman et al., 2022). Our findings identify differential SYT7 abundance as an additional regulator of synaptic diversity between Ib and Is MNs. SYT7 is enriched at the weaker Ib terminals relative to Is synapses that display enhanced release (Fig. 1). Consistent with its inhibitory function, SYT7 loss preferentially enhanced *P_r_* at Ib synapses, with little effect at Is terminals (Fig. 2). In contrast, SYT7 overexpression strongly suppressed release at both synapse types (Fig. 3), indicating endogenous differences in SYT7 abundance contribute to their distinct release properties without reflecting an intrinsic inability of Is terminals to respond to SYT7. Other characteristic Ib and Is MN differences, including increased SV diameter in Is (Fig. 6A, B) and larger synaptic boutons in Ib (Fig. S1K), were not differentially altered in *Syt7* null mutants (Guan et al., 2020) or following SYT7 overexpression (this study). These data indicate SYT7 regulates functional properties rather than determining overall MN identity and morphological features. Beyond the role of SYT7 in lowering SV fusogenicity directly at AZs, our data indicate SYT7 localization to peri-AZs and internal membrane compartments reduces SV availability by promoting and/or stabilizing bulk endosomes that retain SV proteins.

Although we find SYT7 dramatically reduces both spontaneous and evoked release, the number of morphologically docked SVs at active zones is largely preserved (Fig. 6). Immunostaining shows SV proteins like SYT1 and CPX can still cluster at AZs, with only modest changes in their abundance (Fig. 5). Thus, elevated SYT7 can profoundly suppress neurotransmission without altering docked SVs that are positioned for release at AZs. These observations are consistent with reduced SV fusogenicity or impaired functional priming rather than a primary defect in morphological docking. Remarkably, high-frequency stimulation partially overcame SYT7-mediated inhibition, producing robust facilitation in terminals that exhibited minimal release during low-frequency stimulation (Fig. 4). This effect was rapidly reversible, with release returning to strongly suppressed levels following the stimulation train. Prior studies implicated SYT7 as a key Ca²⁺ sensor for short-term synaptic facilitation (Chen et al., 2017; Jackman et al., 2016; Turecek and Regehr, 2018; Weyrer et al., 2021) and in activity-dependent SV docking (Wu et al., 2024). In *Drosophila*, SYT7-dependent suppression of initial SV *P_r_* has been proposed to promote facilitation during repetitive stimulation (Guan et al., 2020; Fujii et al., 2021). Recent structure-function studies suggest that residual Ca²⁺ binding to the SYT7 C2A domain relieves C2B-dependent inhibition of fusion (Sakurai et al., 2026). Although the precise mechanisms underlying facilitation are still unclear, increased release could reflect contributions from activity-dependent modulation of SYT7 inhibition, enhanced residual Ca²⁺ signaling, and the low initial *P_r_* established by SYT7 overexpression.

Although high frequency stimulation in SYT7 overexpression animals enhanced release, facilitation did not restore evoked responses to control levels, indicating substantial inhibition persisted during sustained stimulation. This incomplete recovery suggests elevated SYT7 may influence additional steps in the SV cycle beyond fusion. The observation that SYT7 overexpression also alters presynaptic membrane organization raises the possibility that changes in SV trafficking contribute to the sustained reduction in synaptic output. Consistent with this model, elevated SYT7 caused striking changes in the organization of presynaptic membranes (Fig. 5). SYT1 and CPX partially redistributed from the characteristic halo pattern for SVs near the presynaptic membrane into more irregular internal accumulations, while FM1-43 labeling revealed prominent punctate compartments following intense stimulation. Ultrastructural analysis further demonstrated a substantial increase in enlarged membrane-bound cisternae (Fig. 6). These changes occurred despite relatively preserved SV docking, suggesting elevated SYT7 alters membrane trafficking or the organization of internal SV-associated compartments.

Large presynaptic cisternae can arise through activity-dependent bulk endocytosis, a pathway engaged during intense neuronal activity that retrieves substantial amounts of plasma membrane for subsequent SV regeneration (Cheung and Cousin, 2013). We previously identified enhanced refilling of the RRP in *Syt7* mutants, and suppression of RRP refilling following SYT7 overexpression (Guan et al., 2020). In the current work we find elevated SYT7 levels redistribute SV-associated proteins and promote the accumulation of enlarged presynaptic cisternae, suggesting SYT7 may regulate the bulk endocytosis pathway. Consistent with this model, *Syt7* null mutants exhibited significantly fewer enlarged cisternae following 100 Hz stimulation (Fig. 6). Together, the gain-and loss-of-function phenotypes indicate SYT7 contributes to the regulation of these presynaptic membrane compartments. Whether SYT7 directly promotes bulk membrane internalization, influences cisternae maturation, or alters their subsequent resolution into SVs remains unresolved. The redistribution of Rab5, Rab11, and the endolysosomal marker NHE3 in SYT7-overexpressing terminals suggests broader effects on membrane trafficking within presynaptic terminals (Fig. S2). SYT7 has established functions in endolysosomal trafficking in other cell types (Martinez et al., 2000; Flannery et al., 2010; Czibener et al., 2006), raising the possibility that related activities contribute in part to its presynaptic function. Together, these findings indicate SYT7 regulates synaptic output through effects on both SV fusion and presynaptic membrane trafficking.

The mechanisms establishing differential SYT7 abundance at tonic and phasic synapses remain unknown. Differential SYT7 protein abundance is not accompanied by a corresponding difference in *Syt7* mRNA levels based on Patch-seq analysis of Ib and Is MNs (Jetti et al., 2023), suggesting post-transcriptional regulation contributes to its differential synaptic accumulation. Subtype-specific differences in mRNA translation, protein trafficking, or turnover of SYT7 represent candidate pathways. In mammalian neurons, Presenilin/γ-secretase-dependent processing controls SYT7 protein stability and presynaptic abundance (Barthet et al., 2018; Vevea et al., 2021), while palmitoylation regulates its membrane association and intracellular trafficking (Vevea et al., 2021; Flannery et al., 2010). SYT7 expression can also be regulated post-transcriptionally by microRNAs in non-neuronal cells (Sun et al., 2024). Whether related mechanisms contribute to differential SYT7 abundance at Ib and Is terminals is unknown. A similar mismatch in mRNA and synaptic protein abundance was observed for Tomosyn, another inhibitor of SV fusion enriched at Ib terminals that acts to sequester t-SNAREs (Sauvola et al., 2021). Tomosyn and SYT7 both act as presynaptic release inhibitors and show additive effects in double mutants, indicating they act through genetically separable mechanisms to decrease *P_r_*. Determining how SYT7 and Tomosyn abundance is established and maintained may reveal additional mechanisms through which neurons tune presynaptic output independent of transcriptional differences.

In summary, these findings extend the role of SYT7 beyond its established function in short-term plasticity by identifying a contribution to neuronal subtype-specific synaptic output and presynaptic membrane cycling. Differences in SYT7 expression have also been observed across neuronal subtypes in the mammalian cerebellum (Turecek and Regehr, 2019). In addition, reintroducing SYT7 into climbing fiber synapses that normally lack the protein caused a reduction in initial *P_r_* and a large increase in paired-pulse facilitation (Weyrer et al., 2021). As such, differences in SYT7 levels may represent a more broadly used mechanism for modifying presynaptic output beyond tonic and phasic MN subtypes. Our data indicate SYT7 influences SV availability through two complementary mechanisms in *Drosophila*: (1) regulating the fusion competence of SVs at AZs; and (2) modulating their trafficking through internal membrane compartments. Future studies examining the dynamics and molecular identity of SYT7-associated compartments will be important for defining specific trafficking steps regulated by this Ca²⁺ sensor. Together with the activity of Tomosyn, these findings highlight how the abundance of multiple inhibitory regulators of the SV fusion machinery diversify synaptic output and contribute to tonic– phasic synaptic release differences.

## Materials and methods

### Drosophila stocks

*Drosophila melanogaster* were maintained on standard molasses food at 25°C. OK6-GAL4 (#64199) and *elav*^C155^-GAL4 (#8765) were obtained from the Bloomington Stock Center. The *Syt7*^M1^ null and *Syt7*^GFP^ knock-in were previously described (Guan et al., 2020). CRISPR/Cas9 was used to insert mRuby3 into the endogenous *Bruchpilot* (*Brp)* locus near a previously described MiMIC (MI02987) site (Nagarkar-Jaiswal et al., 2015) to generate BRP^Ruby^. The insertion site was positioned after the BRP sequence RTQAELRV using three *Brp*-targeting gRNAs: GATGACCACGTCCCAGGAAC, TATCCATATCGCCCCAACCG, and TGCACTTGCAATTGCGAGTT. gRNAs and donor constructs were synthesized by Twist Bioscience and assembled into pCFD5 using NEBuilder HiFi DNA Assembly (NEB, E5520). Constructs were injected into *nos-Cas9* embryos (BDSC #78782; BestGene Inc.). For *P_r_* mapping, CRISPR-tagged Cacophony (Cac^RFP^) (Gratz et al., 2019), GluRIIB^GFP^ (provided by Stephan Sigrist) or BRP^Ruby^ were combined with LexAOp-myr-jGCaMP8s (Sauvola et al., 2021) or myr-GCaMP7s as indicated. A 1,251-bp fragment encoding SYT7-RM (FBgn0039900; Twist Bioscience) was cloned into pUAST and injected into *w*^1118^ embryos (BestGene Inc.). Two independent insertion lines, UAS-SYT7^OE1^ and UAS-SYT7^OE2^, were selected based on expression levels.

### Immunocytochemistry and image analysis

3^rd^ instar larvae were dissected in Ca²⁺-free HL3.1 (70 mM NaCl, 5 mM KCl, 4 mM MgCl₂, 10 mM NaHCO₃, 5 mM trehalose, 115 mM sucrose, 5 mM HEPES, pH 7.2), fixed in 4% paraformaldehyde for 10 min, and immunostained as described (Guan et al., 2020). Primary antibodies included mouse anti-BRP (nc82, 1:500; DSHB), rabbit anti-Complexin (CPX, 1:5000) (Huntwork and Littleton, 2007), rabbit anti-SYT1 (1:1000) (Mackler et al., 2002), and rabbit anti-SYT7 (1:500) (Adolfsen et al., 2004). Alexa Fluor 488 goat anti-rabbit (A32731) and Alexa Fluor 555 goat anti-mouse (A21422; Invitrogen) secondary antibodies were used at 1:1,000. Neuronal membranes were labeled with DyLight 649-conjugated anti-HRP (123-605-021) or Cy3-conjugated anti-HRP (123-165-021; Jackson ImmunoResearch) at 1:1,000. NMJs on muscles 1, 4, and 6/7 (segments A2–A4) were imaged using a PerkinElmer CSU-X1 spinning-disk confocal microscope with a Hamamatsu C9100-13 EMCCD camera and 63×/1.40 NA oil objective (Zeiss). Maximum-intensity projections were generated in Volocity 6.5, except for SYT1 and CPX distribution analyses that used single optical sections. Background-subtracted fluorescence was quantified in Fiji (ImageJ2, v2.16.0/1.54p) and normalized to HRP-labeled NMJ area or, for BRP puncta area for AZ localized SYT1 and CPX. Bouton area was measured from manually outlined HRP-positive boutons, and bouton density was calculated per µm of Ib or Is branch length. SYT1 and CPX distributions in MN1-Ib boutons were measured as the ratio of mean fluorescence in the entire HRP-defined bouton (ROI1) to an eroded interior region (ROI2). Fluorescence profiles were additionally obtained from line scans across bouton centers.

### Western blotting

Three adult male heads per biological replicate were homogenized and processed as described previously (Guan et al., 2020). For OK6>SYT^OE2^, a combination of escaper and pharate adult heads were used given the low eclosion rate. Membranes were probed with preadsorbed rabbit anti-SYT7 (1:200) and mouse anti-tubulin (1:2,000,000; Sigma T5168), followed by IRDye 680 goat anti-mouse (926-68020) and IRDye 800CW goat anti-rabbit (926-32211; LI-COR) at 1:20,000. Blots were imaged using a Bio-Rad ChemiDoc system, and band intensities were quantified in Fiji.

### Developmental and behavioral assays

Crosses were maintained at 25°C in five to six replicate vials, with parents removed after 24 hrs. Pupae present on day 7 were marked and monitored daily through day 12 to determine eclosion rates. Pupation height was measured from the bottom of each vial. Locomotor assays were performed as described previously (Nichols et al., 2012). For crawling assays, larvae were placed in 60 mm dishes over graph paper, acclimated for 1 min in HL3.1, and the number of grid lines crossed per min was recorded over two trials.

### FM1-43 dye uptake and unloading

Dissected 3^rd^ instar larvae were incubated with 5 µM FM1-43 (Thermo Fisher, T35356) for 5 min before stimulation. For 10-Hz stimulation, preparations were maintained in HL3 containing 1 mM Ca²⁺ and 20 mM Mg²⁺; for 100-Hz stimulation, HL3.1 contained 1.5 mM Ca²⁺ and 4 mM Mg²⁺. Dye loading was performed at 10 Hz for 5 min or 100 Hz for 30 sec. Following loading, preparations were washed for 1 min with 100 µM ADVASEP-7 (Biotium, #70029) to remove extracellular dye. For unloading, nerves were stimulated continuously at 10 Hz, with fluorescence measured at 1 min intervals and normalized to the initial post-loading intensity. Muscle 4 NMJs in segments A2–A3 were imaged using a Zeiss Axio Imager 2 spinning-disk confocal microscope with a 63× water-immersion objective. Mean NMJ fluorescence was quantified using Volocity 3D Image Analysis software (Quorum Technologies Inc.).

### Electrophysiology and *P_r_* mapping

Two-electrode voltage-clamp (TEVC) recordings were performed from muscle 6 in abdominal segment A3 of male 3^rd^ instar larvae at a holding potential of −80 mV, as previously described (Jorquera et al., 2012). Recordings were performed in HL3.1 (pH 7.2) containing 1.5 mM Ca²⁺ and 4 mM Mg²⁺. Motor nerves were stimulated at the indicated frequencies using a Master-8 programmable stimulator (AMPI). Data were acquired and analyzed using Axoscope 10.0 and Clampfit 10.0 (Molecular Devices), and miniature excitatory junctional currents (mEJCs) were analyzed using Mini Analysis 6.0.3 (Synaptosoft). Inward currents are displayed as positive values. For *P_r_* mapping, 3^rd^ instar larvae were dissected in Ca²⁺-free HL3 containing 20 mM Mg²⁺. Following dissection, preparations were equilibrated for 5 min in HL3 containing 20 mM Mg²⁺ and either 1.0 mM or 1.5 mM Ca²⁺, depending on the experiment. For experiments in *Syt7* null mutants (Fig. 2), 1.0 mM Ca²⁺ was used with myr-GCaMP7s as the fluorescence reporter and endogenously tagged Cac^RFP^ to identify AZs. For SYT7 overexpression experiments using OK6>SYT7OE1 (Fig. 3), 1.5 mM Ca²⁺ was used with myr-GCaMP8s and endogenously tagged BRP^Ruby^. The higher Ca²⁺ concentration improved detection of release events in SYT7-overexpressing terminals with strongly reduced *P_r_*. Muscle 4 NMJs in segments A3–A4 were imaged using a PerkinElmer CSU-X1 spinning-disk confocal microscope with a Hamamatsu C9100-13 EMCCD camera and 60×/1.10 NA water-immersion objective (Olympus). GCaMP fluorescence was recorded at 8 Hz during 1 Hz nerve stimulation for 3–5 mins. At the beginning of each experiment, dual-channel z-stacks of were acquired to identify AZ positions based on Cac^RFP^ or BRP^Ruby^ puncta. Single-plane GCaMP fluorescence recordings were acquired during 1 Hz nerve stimulation delivered through a suction electrode for 3–5 min. A single reference image of AZ positions was acquired every 25 sec to monitor spatial alignment. Minor axial drift was corrected manually, whereas recordings with substantial muscle movement were excluded.

Initial dual-channel z-stacks identified active zones using BRP^Ruby^ puncta, with subsequent imaging every 25 sec to correct for drift. Recordings with substantial muscle movement were excluded. Initial dual-channel z-stacks were converted to maximum-intensity projections using Volocity 6.5.0 (Quorum Technologies Inc.). AZ positions were identified using the Volocity spot-finding function, and equally sized regions of interest (ROIs) were assigned to each AZ. Structures missed by automated detection were annotated manually. GCaMP fluorescence transients were assigned to individual ROIs based on centroid proximity within 0.8 µm. Event times and spatial coordinates were exported to Microsoft Excel. *P_r_* at each AZ was calculated as the number of evoked events per ROI divided by the total number of stimuli.

### Electron microscopy

Larval preparations were fixed in 4% paraformaldehyde for 10 min, washed in HL3, and transferred to 2.5% glutaraldehyde, 4% paraformaldehyde, 90 mM sodium cacodylate (pH 7.2), and 0.2 mM CaCl₂ for 2 hr. Samples were washed twice in 50 mM glycine and three times in 90 mM sodium cacodylate, postfixed for 1 hr in 1% OsO₄ and 1.5% potassium ferrocyanide, and stained with 2% aqueous uranyl acetate for 1 hr. Following graded acetone dehydration, samples were infiltrated with 1:1 LX-112 resin/acetone for 2 hr, followed by fresh resin for 3 hr and polymerization. Ultrathin sections (50–60 nm) were stained with Sato’s lead citrate for 3 min and imaged at 80 kV using an FEI Tecnai Spirit transmission electron microscope at the MIT Koch Institute. For ultrastructural analysis, preparations were examined at rest or immediately following 100 Hz stimulation. SV diameters were measured from electron micrographs, and SVs adjacent to or within 200 nm of the T-bar were quantified. Vesicle diameter distributions were analyzed, with structures larger than 50 nm classified as cisternae.

### Experimental design and statistical analysis

Statistical analyses and graphing were performed using Origin (OriginLab) or GraphPad Prism. Statistical tests are specified in the figure legends. Data are presented as mean ± SEM unless otherwise indicated. Statistical significance is denoted as: *, p≤0.05; **, p≤0.01; ***, p≤0.001 and ****, p≤0.0001.

Supplemental material includes Figures S1 and S2, along with a supplemental data excel file with raw numbers and statical analysis for each data set in the figures.

## Acknowledgements

The authors thank the Bloomington *Drosophila* Stock Center (Indiana University, Bloomington, IN; NIH P40OD018537) and the Developmental Studies Hybridoma Bank (University of Iowa, Iowa City, IA) for providing stocks and antibodies, and Dina Volfson for help generating SYT7 overexpression lines. The authors acknowledge the Koch Institute Microscopy Core Facility at MIT for access to EM instrumentation. This work was supported by the Freedom Together Foundation and NIH grants NS40296, MH104536 and NS117588 to J.T.L. The Koch Institute’s core facilities are supported in part by the National Cancer Institute (P30-CA14051).

## Author contributions

Michiko O. Inouye: conceptualization, data curation, formal analysis, investigation, methodology, visualization, and writing-original draft, review, and editing. Yulia Akbergenova: data curation, formal analysis, investigation, and writing-review and editing. Zhuo Guan: data curation, formal analysis, and writing-review and editing. Andrés Crane: data curation, formal analysis, and writing-review and editing. Mónica C. Quiñones-Frías: conceptualization and writing-review and editing. J. Troy Littleton: conceptualization, supervision, project administration, and writing, review and editing-original draft.

---

**Figure S1.**
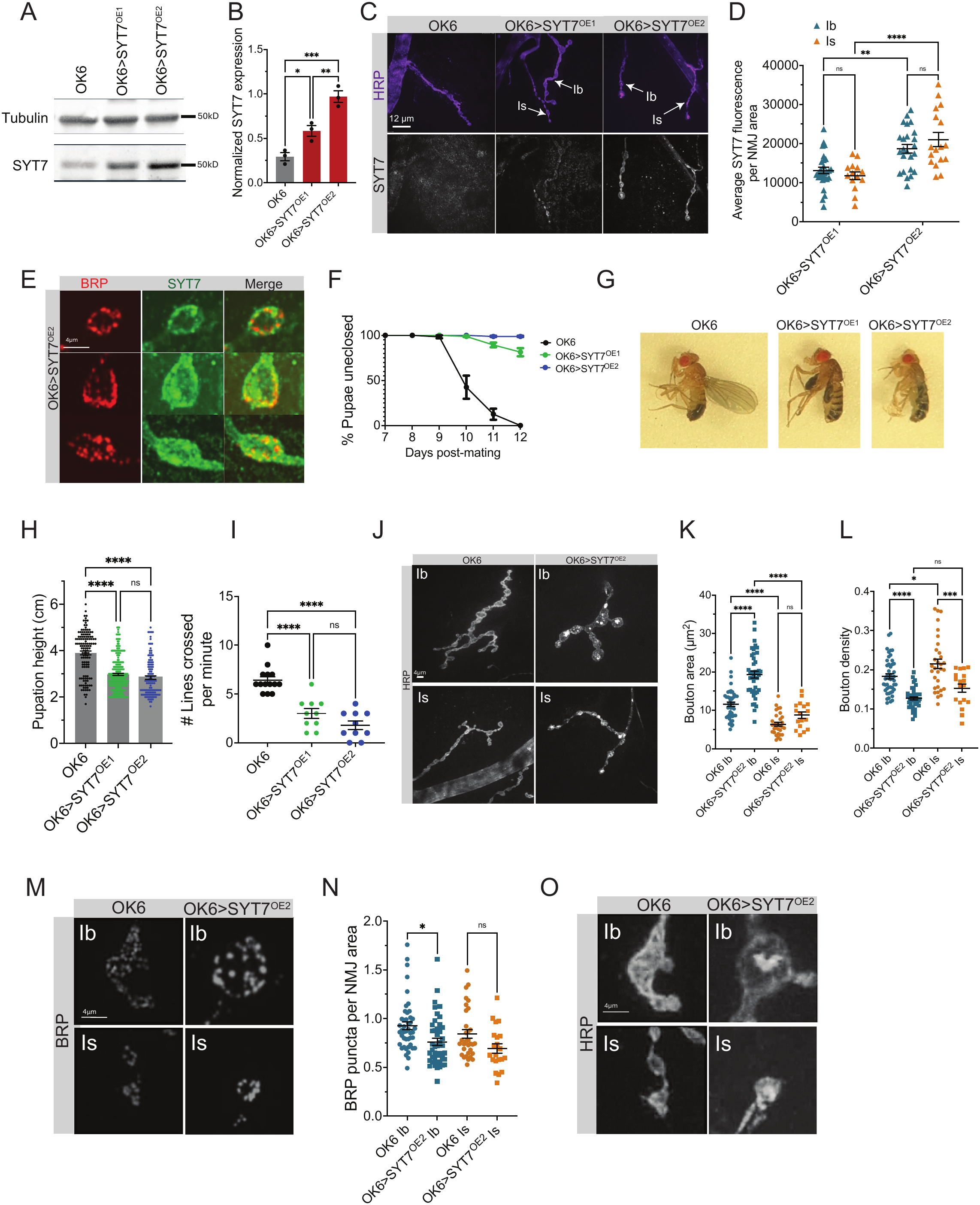
MN-specific SYT7 overexpression results in adult lethality and defective behavior. **(A)** Western blot of adult head extracts from OK6-Gal4, OK6>SYT7^OE1^ and OK6>SYT7^OE2^, probed with anti-SYT7 and anti-Tubulin (loading control). Scale bar = 12 μm**. (B)** Quantification of normalized SYT7 to Tubulin expression from western analysis. **(C)** Immunocytochemistry of 3^rd^ instar muscle 4 NMJs with anti-HRP (magenta) and anti-SYT7 (grey) in OK6-Gal4, OK6>SYT7^OE1^ and OK6>SYT7^OE2^ larvae. **(D)** Quantification average SYT7 fluorescence per NMJ area for the indicated genotypes. **(E)** Representative Ib bouton single-plane confocal images of 3^rd^ instar larval muscle 4 NMJs immunostained with anti-SYT7 (green) and anti-BRP (red) in OK6>SYT7^OE2^ larvae. MN-specific overexpression of SYT7 leads to reduced eclosion rate **(F)**, unexpanded wings **(G)**, and decreased mobility in larvae as indicated by reduced pupation height **(H)** and larval crawling rate per minute **(I)**. **(J)** Representative images of HRP immunolabeling within Ib and Is synaptic boutons of 3^rd^ instar M4 NMJs of controls and OK6>SYT7^OE2^ larvae, with quantifications for bouton area **(K)** and bouton density **(L)**. **(M, O)** Images of BRP and HRP immunolabeling, with quantification of BRP puncta number normalized to HRP area **(N)**. Data = mean ± SEM. Statistical analysis was performed with one-way ANOVA with Tukey correction. * p<0.05; *** p<0.001; **** p<0.0001; ns, not significant.

**Figure S2.**
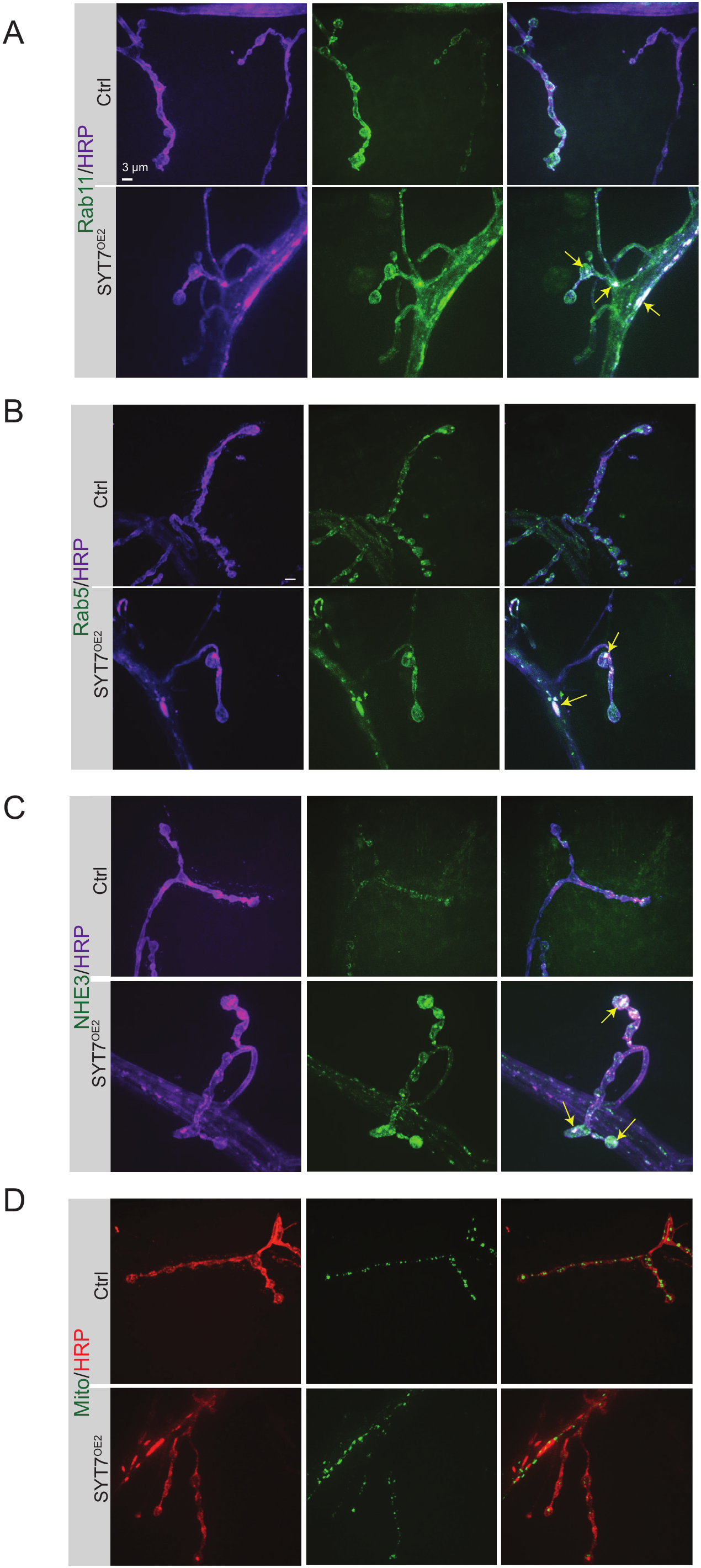
SYT7 overexpression leads to large presynaptic internal compartments. Anti-HRP immunostaining in controls and SYT7-overexpressed 3^rd^ instar larval NMJs immunostained with anti-HRP and the late endosomal marker Rab11 **(A)**, the early endosomal marker Rab 5 **(B)**, the lysosomal marker NHE3 **(C)** and mitochondria **(D)**. For panel A, control is elav^C155^-Gal4; UAS-Rab11-GFP and SYT7^OE2^ is elav^C155^-Gal4; UAS-SYT7^OE2^/UAS-Rab11-GFP. For panel B, control is elav^C155^-Gal4;; UAS-Rab5-GFP and SYT7^OE2^ is elav^C155^-Gal4; UAS-SYT7^OE2^; UAS-Rab5-GFP. For panel C, the control is elav^C155^-Gal4;;UAS-NHE3-GFP and SYT7^OE2^ is elav^C155^-Gal4; UAS-SYT7^OE2^; UAS-NHE3-GFP. For panel D, control is elav^C155^-Gal4;; UAS-Mito-GFP and SYT7^OE2^ is elav^C155^-Gal4; UAS-SYT7^OE2^; UAS-Mito-GFP. Arrows point to presynaptic compartments containing endosomal and lysosomal, but not mitochondrial markers, that co-stain with HRP. Scale bar = 3 μm.

